# Scalable *In Vitro* Production of Defined Mouse Erythroblasts

**DOI:** 10.1101/2020.11.10.376749

**Authors:** Helena S Francis, Caroline L Harold, Robert A Beagrie, Andrew J King, Matthew E Gosden, Joseph W Blayney, Danuta M Jeziorska, Christian Babbs, Douglas R Higgs, Mira T Kassouf

## Abstract

Mouse embryonic stem cells (mESCs) can be manipulated *in vitro* to recapitulate the process of erythropoiesis, during which multipotent cells undergo lineage specification, differentiation and maturation to produce erythroid cells. Although useful for identifying specific progenitors and precursors, this system has not been fully exploited as a source of cells to analyse erythropoiesis. Here, we establish a protocol in which characterised erythroblasts can be isolated in a scalable manner from differentiated embryoid bodies (EBs). Using transcriptional and epigenetic analysis, we demonstrate that this system faithfully recapitulates normal primitive erythropoiesis and fully reproduces the effects of natural and engineered mutations seen in primary cells obtained from mouse models. We anticipate this system to be of great value in reducing the time and costs of generating and maintaining mouse lines in a number of research scenarios.

**Key Points:**

- Scalable purification of primitive-like erythroid cells from *in vitro* differentiated mESCs offers tractable tools for genetic studies
- *In vitro* derived erythroid cells recapitulate wild type and engineered mutation phenotypes observed in primary cells obtained from mouse models

## INTRODUCTION

The isolation of embryonic stem cells from developing mouse blastocysts, their maintenance in culture, and genetic manipulation has provided a fundamentally important research tool for experimental biology.^1^ *In vitro* differentiation of stem cells offers unparalleled access to developmental pathways including well defined multipotent cells, precursors and mature cell types representing a wide range of organ systems.^2^ Developing robust protocols using mESCs to obtain specific cell types at scale would further allow the use of these cells for detailed molecular analysis and large scale, high throughput screens. Nevertheless, for maximum value, it is crucial that mESC-derived cells and tissues faithfully represent the corresponding primary cell populations.

During development, mouse haematopoiesis occurs in three distinct waves. Primitive haematopoiesis originates in the blood islands of the yolk sac (embryonic day E7.25-8.5). This is transiently accompanied by definitive haematopoiesis arising from Erythroid Myeloid Progenitors (EMP; E8.25-9.5) and eventually replaced by long-term definitive haematopoiesis emerging from the aorta-gonad-mesonephros (AGM) at around E10.5.^3–5^ Lineage specification, differentiation and maturation of haematopoietic cells at each stage of development has been extensively investigated using mESCs.^2,3,6^ Such studies have informed researchers of normal blood formation and also elucidated some of the mechanisms underlying inherited and acquired blood diseases.^3,5,7^ To investigate haematopoietic differentiation, mESCs are cultured in the absence of leukaemia inhibitory factor (LIF), which results in the formation of a 3D organoid or embryoid body (EB). This promotes differentiation that faithfully represents early mouse development; resulting EBs therefore contain a mixture of all three germ layers.^8^ The use of additional cytokines, alternative plating strategies, and/or re-plating can be used to support particular differentiation pathways of interest.^6^

The precise origins and cellular outputs of mESC-EB cultures have been of main interest to the field of developmental biology for years.^4,6,9,10^ In hematopoiesis, initial studies used colony re-plating assays of EB-derived cells and detailed the emergence of erythroid progenitors of primitive and definitive nature from day four onwards.^11^ This was subsequently shown to closely reflect erythropoiesis in mouse development *in vivo*.^3,12^ Within a similar time-frame, haemoglobinized erythroid cells also begin to arise in cultured EBs.^11^ Whilst in early studies haemoglobinization was observed in only ~1% of EBs, optimisation of culture conditions has now increased erythrocyte-containing EB production to almost 100%.^13,14^ Isoelectric focusing,^13^ RNase protection,^15^ and RT-PCR data^11^ from earlier studies identified embryonic globins, suggesting the presence of erythrocytes that most likely recapitulate some aspects of primitive erythropoiesis. Later studies showed that under specific conditions, the cultures may yield both primitive and definitive AGM-like progenitor cells.^10^ More recent immunophenotypic characterization and re-plating of progenitors revealed that the primitive and definitive outputs detected in mESC-EB culture resemble primitive and EMP outputs in mouse embryos.^4^ It is not yet clear if truly definitive long-term repopulating hematopoietic stem cells are represented in these cultures.^10,16–17^ As demonstrated over almost two decades of work in this system, the co-emergence of progenitors of mixed origins hinders the use of the culture system as a source of lineage-specific haematopoietic cells for detailed molecular studies and obviate the need for further characterisation of the cells produced. If better defined, the erythroid cell production in the mESC-EB system would expand the *in vitro* system from a developmental biology platform to a mammalian genetics and molecular screening platform.

In this study, we exploit the spontaneous differentiation of erythroid cells within EBs as a readily-accessible and scalable erythroid cell population. By selecting erythroid cells expressing the transferrin receptor (CD71), we can isolate and characterize a large population of mouse erythroid cells. We show that this population is homogeneous, faithfully represents normal primitive erythropoiesis and accurately mimics the resulting phenotypic effects of genetic manipulations seen in primary cells obtained from mice harbouring the same genetic perturbations, potentially avoiding the need to establish full mouse models to assess the effects of such manipulations. Finally, we present a protocol for miniaturization of the differentiation protocol, which offers the potential to perform high-throughput studies on tens to hundreds of genetic models in a single experiment.

## METHODS

### Mouse embryonic stem cell culture and genome engineering

E14TG2a.IV mESCs were maintained by standard methods.^18,19^ To produce genetically modified mESC models, variations of CRISPR/Cas9 strategies were used in conjunction with homology-directed repair (HDR) when needed (YFP-tagged α-globin and D3839 models). mESCs were co-transfected with guide RNA and HDR vectors using TransIT-LT1 reagent (Mirus; according to manufacturer’s instructions) for YFP-tagged α globin and Neon electroporation system (Invitrogen, according to manufacturer’s instructions) for DelR1 and D3839 mutants. Details on transfection conditions and sequences for guide RNA (Supplemental Table 1) and HDR vectors are in Supplemental Methods.

### EB differentiation and erythroid population isolation

24h prior to differentiation, mESCs were induced by passaging into IMDM based media supplemented with LIF. To start the differentiation culture (d0), cells growing in IMDM were trypsinized and plated in differentiation media in either triple vent petri dishes (Thermo Fisher) or flat-bottom 96-well plates (Thermo Fisher) at 3×10^4^ cells in 10 cm dishes or 100-1200 cells per well of a 96-well plate for up to seven days without further intervention except for daily gentle shaking of the dishes to disrupt potential EB attachment to the bottom of the dish. EBs were harvested and disaggregated in 0.25% trypsin for 3 minutes at 37°C. See Supplemental Methods for details.

CD71+ cells were isolated by magnetic column separation (LS Column, Miltenyi), according to the manufacturer’s instructions. Briefly, cells from disaggregated EBs were labelled with anti-mouse CD71-FITC (eBioscience 11-0711-85; 1:200) in staining buffer (PBS with 10% FCS; 500 μl per 10^7^ cells) for 20 minutes at 4°C, washed, then incubated with MACS anti-FITC separation microbeads (Miltenyi; 10 μl per 10^7^ cells, according to manufacturer’s instructions). Bead-labelled cells were retained by LS columns. See Supplemental Methods for details.

### Flow cytometry

Cells were stained with antibodies for 20 minutes at 4°C in staining buffer. Stained cells were analysed using an Invitrogen Attune NxT cytometer (Thermo Fisher) and the FlowJo software package (BD). See Supplemental Methods for details.

### ATAC-seq and ChIP-seq

ATAC-seq was performed on 70×10^3^ cells from target populations as previously described.^20,21^ ChIP-seq was performed as described^22^ on aliquots of 5×10^6^ CD71+ cells derived from two separate differentiation experiments per antibody. Chromatin fragmentation was performed for 8 minutes as optimized for the use of Bioruptor Pico sonication device. Data from both methods were analysed with an in-house pipeline as described.^21,23^ In each case, for visualization, alignment files from two or three biological replicates were normalized to reads per kilobase per million mapped reads (RPKM) and averaged.

To compare datasets genome-wide, triplicate data for each tissue of interest were peak-called with the Model-based Analysis of ChIP-Seq tool (MACS2)^24^ using default parameters. PCA was then performed using the DiffBind package in R.^25^

### Next-generation Capture-C

Next-generation Capture-C was performed as previously described^26^ on 5×10^6^ mouse CD71+ cells derived from three separate differentiation experiments. To visualize differences in Capture-C profiles, normalized interactions from D3839 erythroid cells were subtracted from wild-type erythroid interactions to generate a differential Capture-C track.

### RNA expression analysis

RT-PCR: 10^6^ cells were lysed in TRI reagent (Sigma) and RNA extracted using Direct-zol columns with a 30 minute on-column DNase step at room temperature (Zymo Research). cDNA was generated with Superscript III First-Strand Synthesis SuperMix (Life Technologies). For primers, see Supplemental Table 2.

Single-cell RT-PCR: single-cell expression analysis necessitated isolation of 185 single erythroid cells by indexed FACS into 96-well plates. See Supplemental Methods for details. Briefly, cells were lysed, RNA reverse transcribed and material used for 43 TaqMan assays (Supplemental Table 3 and 4). When performing reverse transcription and pre-amplification on sorted single cells, one aliquot of RNA standard was included (prepared and used as detailed in supplemental data).

Data analysis was performed using Python (v3.5.2; Matplotlib v2.0.2; Numpy v1.14.5; Pandas v0.23.3; scikit-learn v0.19.2; Scipy v1.1.0). See Supplemental Methods for details.

### Data Sharing statement

ATAC-Seq, ChIP-Seq, and Capture-C data reported in this article have been deposited in the Gene Expression Omnibus database (accession number XXX).

## RESULTS

### A pure erythroid population can be isolated from a 7-day embryoid body culture

The generation of EBs from mESCs is well-established.^6^ To enrich for haematopoietic lineages, cells are plated at low density in differentiation media using bacterial dishes and harvested between days 2 and 7. Allowing EBs to differentiate undisturbed results in high levels of spontaneous erythroblast differentiation and haemoglobinization, observed as red EBs.^14^

To optimize the number and purity of the obtained erythroid cells, we first established the kinetics of erythroid differentiation in the E14 mESC line. As EB growth and disaggregation beyond day 7 of differentiation compromised viability, we focused our investigation on timepoints up to and including day 7. Previously published data show that globin expression increases in EBs as they differentiate in culture.^13,15,27^ We confirmed this by RT-PCR at days 4-7 of differentiation (Figure 1A). Primary erythroid cells are usually staged by immunophenotyping using the erythroid markers CD71 and Ter119, as early erythroid progenitors are marked by CD71 alone and subsequent populations first gain Ter119 and then lose CD71. We examined the expression of these same markers in EBs and confirmed the red cell expansion which occurs in parallel with increased expression of erythroid-specific genes (Figure 1B).^28,29^ By day 7, cells from disaggregated EBs were ~40% positive for CD71, and about 30% also positive for Ter119.

**Figure 1:**
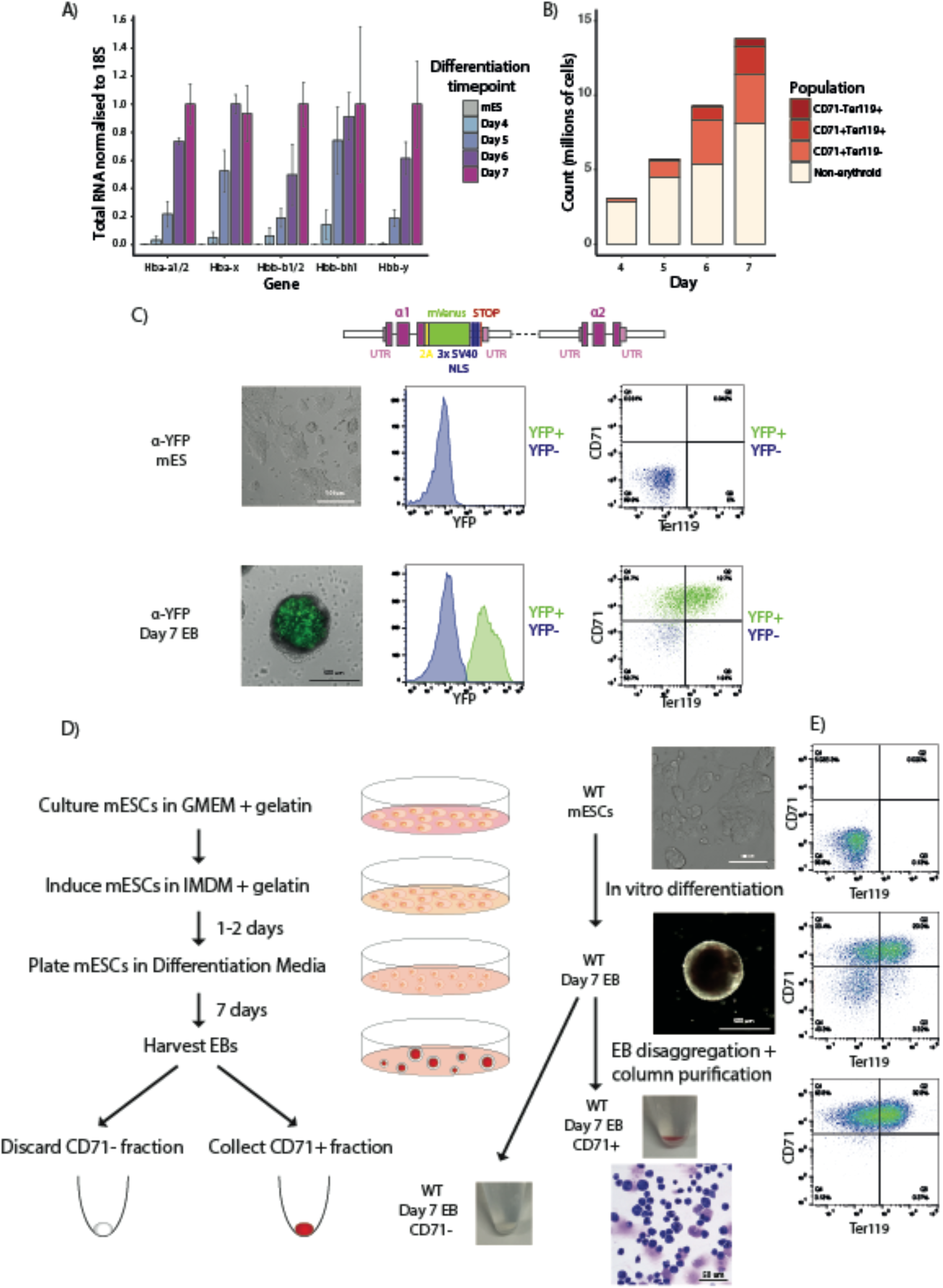
*in vitro* differentiated EBs as a source of erythroid cells. **A**) RT-PCR of mature mouse globin transcripts in mESCs and EBs between days 4-7 of differentiation, normalized to the 18S housekeeping gene. Levels are shown relative to the maximal detected expression for each gene. Bars represent mean values from three independent differentiations; error bars represent standard deviation of the mean. **B**) Cell counts for immunophenotypically-defined populations using antibodies for CD71 and Ter119 cell surface markers through days 4-7 of EB differentiation. Data shown are from a representative differentiation in 10 cm dish format. **C**) Fluorescence levels of an α-globin-YFP tag in mESCs and day 7 EBs. Top: a schematic of the tag shows the insertion site after the final exon of the gene with a 2A self-cleaving peptide sequence in yellow, the mVenus coding region in green, nuclear localization signal (NLS) repeats in blue, the STOP codon in red and the untranslated regions (UTRs) in pink. Bottom: brightfield images of mESCs and a single EB are overlaid with YFP fluorescence signal (left panels). Flow cytometry histograms for YFP fluorescence demonstrate the presence of an α-globin-positive population (green peak) in day 7 EBs (middle panels). Flow cytometry plots for the erythroid markers CD71 and Ter119 show the overlap of YFP+ population from the histogram with the CD71+ cell populations (YFP+ cells labelled green as in the histogram) in day 7 EBs (right panels). **D**) Protocol summary for the generation of EB-derived CD71+ erythrocytes *in vitro*. Example data for column-based CD71+ cell purification, starting with brightfield images of cultured mESCs (top panel), to whole EB (middle panel), to CD71-separated populations shown as cell pellet images for CD71+ (red pellet) and CD71- (clear pellet) fractions. A stained (modified Wright stain) cytospin preparation is shown for the purified CD71+ erythroid population (bottom panel). **E**) Flow cytometry plots for with CD71 and Ter119 markers are shown for populations at each step.

To purify erythroid cells from day 7 EBs, we focused on the transferrin receptor surface protein CD71: a reliable marker of primitive and definitive erythroid cells.^28,29^ To ensure that CD71 identified only erythroid cells from a heterogenous mix of EB cell populations, we monitored globin expression using an mESC line with one copy of the red cell specific α-globin gene heterozygously tagged with a yellow fluorescent protein (YFP). By flow cytometry, all YFP+ cells were also stained for CD71 (Figure 1C) confirming that CD71 is also a robust marker of erythroid cells derived from whole EBs.

Next, we used a magnetic column-based selection method to isolate CD71+ erythroid cells from day 7 EBs (Figure 1D). Compared to FACS-based sorting, the use of a column-based selection results in low levels of cell death and allows for rapid selection of large numbers of cells. Cytospin staining confirmed that the isolated CD71+ cells are a mix of erythroid cell differentiation stages, in agreement with immunophenotyping data (Figure 1D). Moreover, the high stringency magnetic column selection resulted in a high purity, as assessed by flow cytometry (>98% CD71+; Figure 1E), although there was some cell loss; the overall fraction of recovered CD71+ cells was lower (20-25% of the total EB cell count) than the expected 40% CD71+ cells detected in whole EBs. The total cell yield from a typical single 10 cm dish of cultured EBs is 5-10×10^6^ cells (see Methods); we obtain typically around 1-2 x10^6^ CD71+ cells per plate. The cell numbers required for most molecular assays can be attained by scaling appropriately. In conclusion, we have developed a simple, scalable protocol for isolating a pure erythroid cell population from a single plating.

### Erythroid cells from EB cultures are uniformly derived from the primitive lineage

To determine the nature of the EB-derived CD71+ erythroid cells, we next compared DNA accessibility (determined by ATAC-seq) of the EB-derived CD71+ erythroid population with both primitive and definitive primary erythroid cells.^20^ As demonstrated by the DNA accessibility profile of embryonic-specific genes at the globin loci, results from EB-derived cells most closely resemble primitive erythroblasts (Figure 2A). By performing principal component analysis (PCA) of all ATAC-seq peaks called in each dataset, we found that EB-derived cells clustered with primary primitive erythroid cells (E10.5 blood) at a genome-wide level (Figure 2B; Supplemental Figure 1A). This clustering persisted with no discernible pattern associated with red cell differentiation staging; ATAC-seq data from E9.5, E10.5 and E11.5, representing early to more mature primitive erythroid cells respectively^12,30,31^ cluster with EB-derived populations separated based on the CD71/Ter119 markers from less mature (CD71+ only, S1) to more mature erythroid cells (CD71+/Ter119+ double positive, S3) (Supplemental Figure 1B-C).^28^

**Figure 2:**
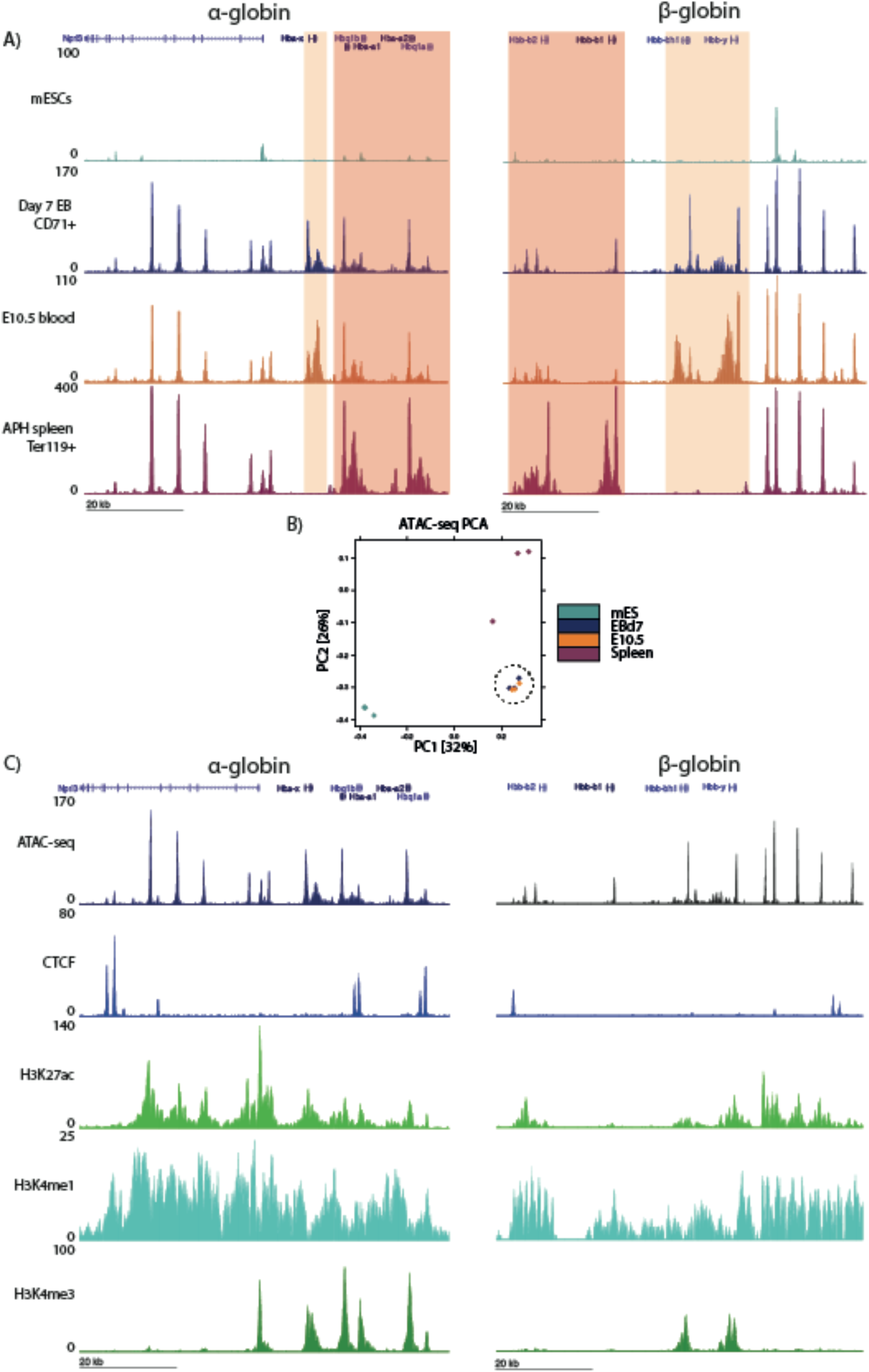
The EB-derived erythroid population resembles embryonic primary red cells by chromatin assays. **A**) ATAC-seq tracks for the α- and β-globin loci (chr11:32,131,800-32,204,799 and chr7:110,952,000-111,027,999, respectively) from mESCs and erythroid cells from day 7 EBs, E10.5 embryonic blood and Ter119+ erythrocytes from adult APH-treated spleen. Tracks are RPKM-normalized and averaged over three biological replicates. Embryo-specific globin genes are highlighted in orange; adult globin genes are highlighted in pink. Other visible peaks represent erythroid-specific enhancer elements. **B**) PCA plot of genome-wide ATAC-seq peaks from the tissues shown above. EB-derived cells cluster with E10.5 primary erythroid cells (highlighted by the dashed circle), away from mESCs and spleen-derived erythrocytes. **C**) CMP-seq tracks for CTCF-ChIP seq and histone marks (H3K27ac and H3K4me1 for enhancers and H3K4me3 for promoters) in EB-derived erythrocytes at the α-and β-globin loci. ChIP-seq data are shown as average tracks across two biological replicates, normalized by RPKM.

To complete the characterization of the chromatin state of EB-derived erythroid cells, we performed ChIP-seq for CTCF and a number of key histone modifications that typically mark active regulatory elements (Figure 2C). The data closely recapitulates the patterns observed in primary embryonic erythroid cell data (A.K. and Duantida Songdej, manuscript submitted September 2020). Together, the chromatin profiling data indicate that the CD71+ EB fraction represents a primitive-like erythroid state at the population level.

Finally, to confirm that EB-derived CD71+ cells represent a homogeneous population of primitive-like erythroid cells, we used single-cell RT-PCR to compare expression of a panel of 40 genes (Supplemental Tables 3 and 4) previously shown to discriminate between the primitive and definitive erythroid lineages across a range of differentiation stages (Supplemental Tables 3 and 4.^31,32^ For primitive primary erythroid cells, we collected blood from embryonic days E9.5, E10.5 and E11.5, representing semi-synchronously differentiating primitive red cells in circulation.^12,29–31^ For definitive erythroid cells, we used established CD71/Ter119-based immunophenotyping and isolated S1-S3 gates^28^ from mouse fetal liver and adult acetylphenylhydrazine-treated (APH) spleen, representing an early (S1) and later (S3) stage of definitive erythroid maturation. For EB-derived erythroid populations, we isolated CD71+ cells which were either positive (S3) or negative (S1) for Ter119 (Supplemental Figure 2A). Focusing on the globin genes, we found that all primary cells showed expected patterns of expression: adult *Hba-a*1/2 genes were expressed in both primitive and definitive cells, *Hbb-b1/2* genes were exclusively transcribed in definitive cells, whereas embryonic *Hba-x* and *Hbb-y* genes were only detected in primitive cells (Figure 3A). All globin genes showed increased expression with cell maturation (Figure 3A). Importantly, all EB-derived cells were simultaneously positive for both embryonic and adult alpha globin expression, demonstrating that the CD71+ EB population is uniformly comprised of the primitive lineage (Figure 3A). By relative expression levels, EB-derived cells most closely resembled E9.5 cells, suggesting that they represent an early stage of erythroid differentiation.

**Figure 3:**
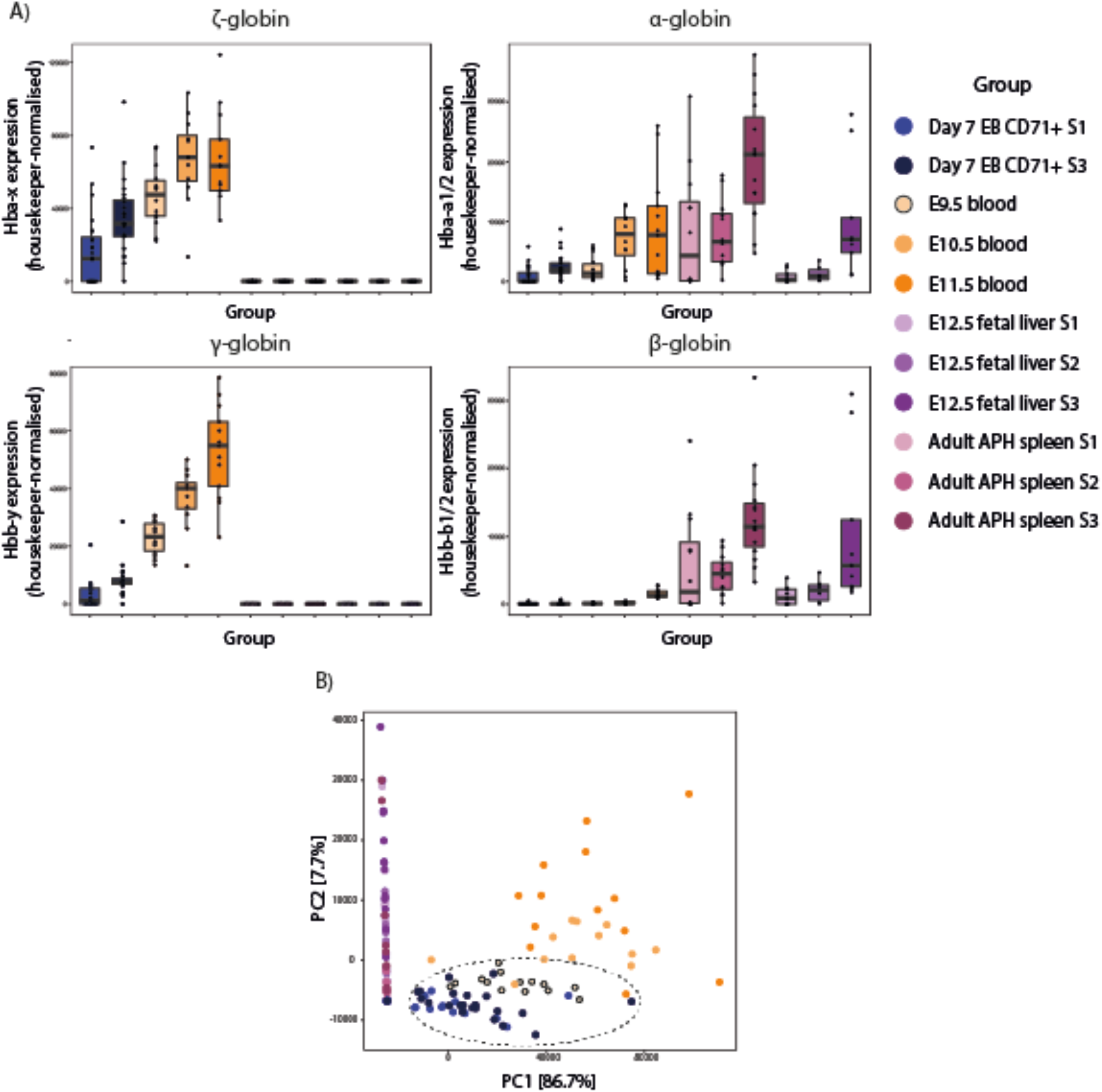
EB-derived erythrocytes are uniformly of the primitive lineage. **A**) Single-cell RT-PCR (Biomark) from FACS-sorted primary and EB-derived erythroid cells as indicated in the colour coded groups for embryonic (*Hba-x* and *Hbb-y*) and adult (*Hba-a1/2* and *Hbb-b1/2*) globin genes. **B**) PCA of expression data from a panel of 40 probes used for single-cell RT-PCR (Biomark) to distinguish primitive and definitive lineages of specific differentiation stages. EB-derived erythroid cells (dark and light blue dots) most closely resemble E9.5 primary cells (orange circles with a black outline) as indicated with the black dotted circle.

The same trend was true using PCA across all gene expression data (gene list in Supplemental Tables 3 and 4): all EB-derived erythroid cells clustered with primitive erythroid cells (particularly with E9.5), and were distinct from definitive cells (Figure 3B). Data for representative individual probes confirmed this finding with expression greatly enriched in either primitive or definitive tissues, as expected (Supplemental Figure 2B). Furthermore, PCA analysis excluding the globin gene data still distinguished between the major populations (Supplemental Figure 2C). Although representing a subset of the transcriptome, the molecular signature presented in this expression dataset provides strong evidence that the spontaneously differentiated erythroid output of EB day 7 is uniform and represents the primitive erythroid pathway.

### Genetically modified EB-derived erythroid cells recapitulate the phenotype of their in vivo derived counterparts

When substituting *in vivo* models with *in vitro* cell systems, one must have confidence that the latter faithfully recapitulates the former.^3,33,34^ We therefore compared the phenotypes of specific genetic manipulations in EB-derived cells with their counterparts in primary cells. We used the α-globin gene cluster as a well characterized model of mammalian gene expression. The α-globin genes are expressed and similarly regulated in both embryonic and adult red cells (A.K. and Duantida Songdej, manuscript submitted September 2020). We initially showed that the pattern of α-globin-like gene expression in the EB-derived erythroid cells closely resembles that seen in normal *in vivo* primitive erythropoiesis (Figure 3A).

To test whether key regulatory elements in this multi-gene locus acted similarly in EB-derived erythroid cells and primary *in vivo* mouse erythroid cells, we analysed the molecular phenotype of erythroid cells derived from genetically modified mESCs and their corresponding *in vivo* mouse models. Deletion of one of the major enhancers of the adult α-globin genes, R1, reduces the expression of alpha globin by 40% in definitive erythroid cells;^21^ the same effect is seen in E10.5 primitive red cells with no associated effect on ζ-globin (A.K. and Duantida Songdej, manuscript submitted September 2020). R1 was deleted from both alleles in ΔR1 mESCs and EB-derived erythroid cells were isolated and analysed for gene expression. ΔR1 EB-derived erythroid cells show downregulation of α-globin as observed in the equivalent mouse model both in E10.5 primitive and fetal liver definitive erythroid cells (Figure 4A). No effect on ζ-globin is observed in ΔR1 EB-derived red cells as in primary E10.5 erythroid cells (Figure 4A; A.K. and Duantida Songdej, manuscript submitted September 2020).

**Figure 4:**
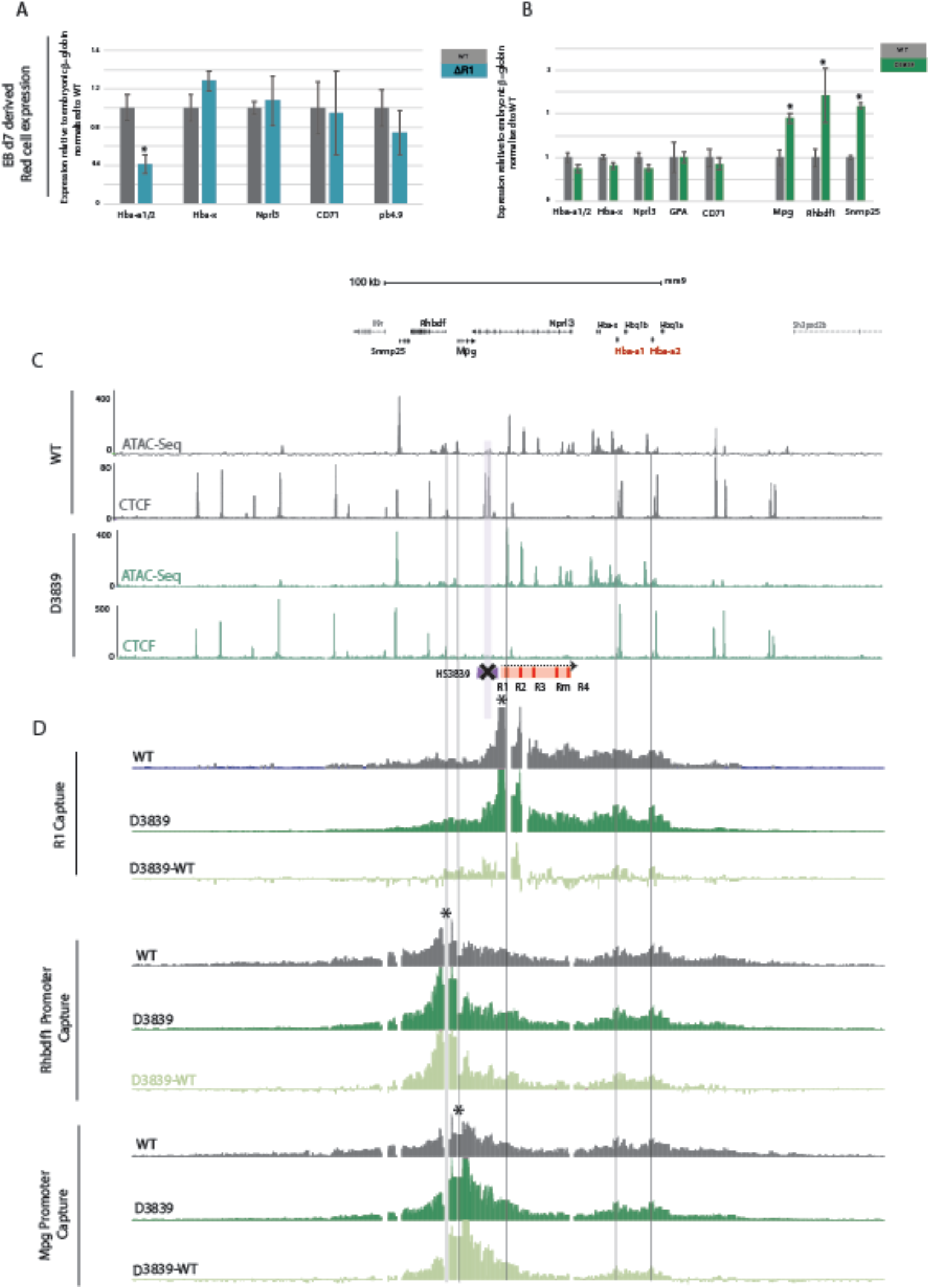
**A**) Expression data for indicated genes based on mature transcripts from enhancer R1 knock out mESCs (ΔR1) day 7 EB-derived erythroid cells, normalized to the embryonic β-globin genes. Levels are shown relative to wildtype day 7 EB-derived erythroid cells (WT). Bars represent mean values from at least six independent differentiations; error bars represent standard deviation of the mean. Student’s *t*-test \**P* <0.001. **B**) Expression data for indicated genes based on mature transcripts from CTCF (HS3839) knock out mESCs (D3839) day 7 EB-derived erythroid cells, normalized to the embryonic β-globin genes. Levels are shown relative to equivalent wildtype cells (WT). Bars represent mean values from at least six independent differentiations; error bars represent standard deviation of the mean. Student’s *t*-test \**P* <0.001. **C**) RPKM-normalized ATAC-seq and CTCF ChIP-seq tracks averaged for three replicates of wildtype and D3839 erythroid cells, both derived from day 7 EBs. **D**) Differential interactions (by NG-Capture-C) of α-globin regulatory regions and flanking genes between WT and D3839 d7 EB-derived erythroid cells. Capture-C data for the indicated viewpoints (black asterisks) in WT and D3839 erythroid cells are shown. Data representing at least 3 independent differentiation for two independently generated clones were used. Differential tracks show a subtraction (D3839-WT) of the mean number of normalized meaningful interactions per restriction fragment.

We also created an mESC line in which the CTCF boundary (HS38-39) was deleted in homozygosity (D3839) from the α-globin cluster as confirmed by ATAC-Seq and CTCF ChIP-Seq (Figure 4C) and compared the phenotype of the EB-derived erythroid D3839 cells to those from the corresponding mouse model.^22^ Again we found that the *in vitro* mESC-EB culture system largely recapitulated the *in vivo* phenotype. When compared to WT, D38-39 EB-derived erythroid cells showed perturbation of gene expression similar to that reported in the D38-39 mouse model;^22^ the *Mpg, Rhbdf1*, and *Snrnp25* genes, located 5’ of the deleted boundary, were upregulated (Figure 4B). Furthermore, an extension of the enhancer/promoter interaction domain to include promoters of the perturbed genes was also revealed by Capture-C in D3839 EB-derived erythroid cells (Figure 4D) as described in the D3839 mouse model.^22^ The perturbed chromatin interaction profile is captured both from the α-globin R1 enhancer as well as the promoters of the affected genes (*Rhbdf1* and *Mpg*); the subtraction tracks (D3839-WT) indicate a gain in significant chromatin interactions between *Mpg, Rhbdf1* promoters and the α-globin cluster (Figure 4D). Recapitulating the complex phenotype of the HS38-39 boundary deletion supports the argument for the mESC-EB system as a faithful *in vitro* model for dissecting complex molecular mechanisms and address current outstanding questions such as the relationship between genome structure and function.

### EB differentiation can be miniaturized to a 96-well format

Rather than calling for mid-to-large numbers of cells of a single genotype, many experimental designs will instead require simultaneous *in vitro* differentiation of multiple mES lines, at a manageable and affordable scale. This is particularly the case for screening-style assays employing large numbers of genetic manipulations, or for testing many small molecules in a high-throughput manner.

To make mESC differentiation scalable for a broader range of applications, we optimized conditions for EB cultures using a 96-well plate format (Figure 5A). By plating a range of cell numbers (100-1200) into 200 μl differentiation media in individual wells of uncoated plasticware, we found that EBs could be formed as seen for bulk cultures; EB-derived erythroid cells from the miniaturized protocol are comparable to those obtained using 10 cm dish format, as judged by flow cytometry and cell morphology (Figure 5B). In addition to the 96-well plate manufacturer, the initial plating density of mESCs was critical for the characteristics of EB formation: higher cell densities at the time of plating resulted in the formation of fewer, larger EBs (Supplemental Figure 3A). The differences in EB aggregation properties had consequences on erythropoiesis: the percentage of CD71+ erythroid cells decreased with increasing EB size (Figure 5C, Supplemental Figure 3C). We found the optimal plating density for maximal CD71+ erythroid cell output is between 200 to 400 mESCs per well (Figure 5C).

**Figure 5:**
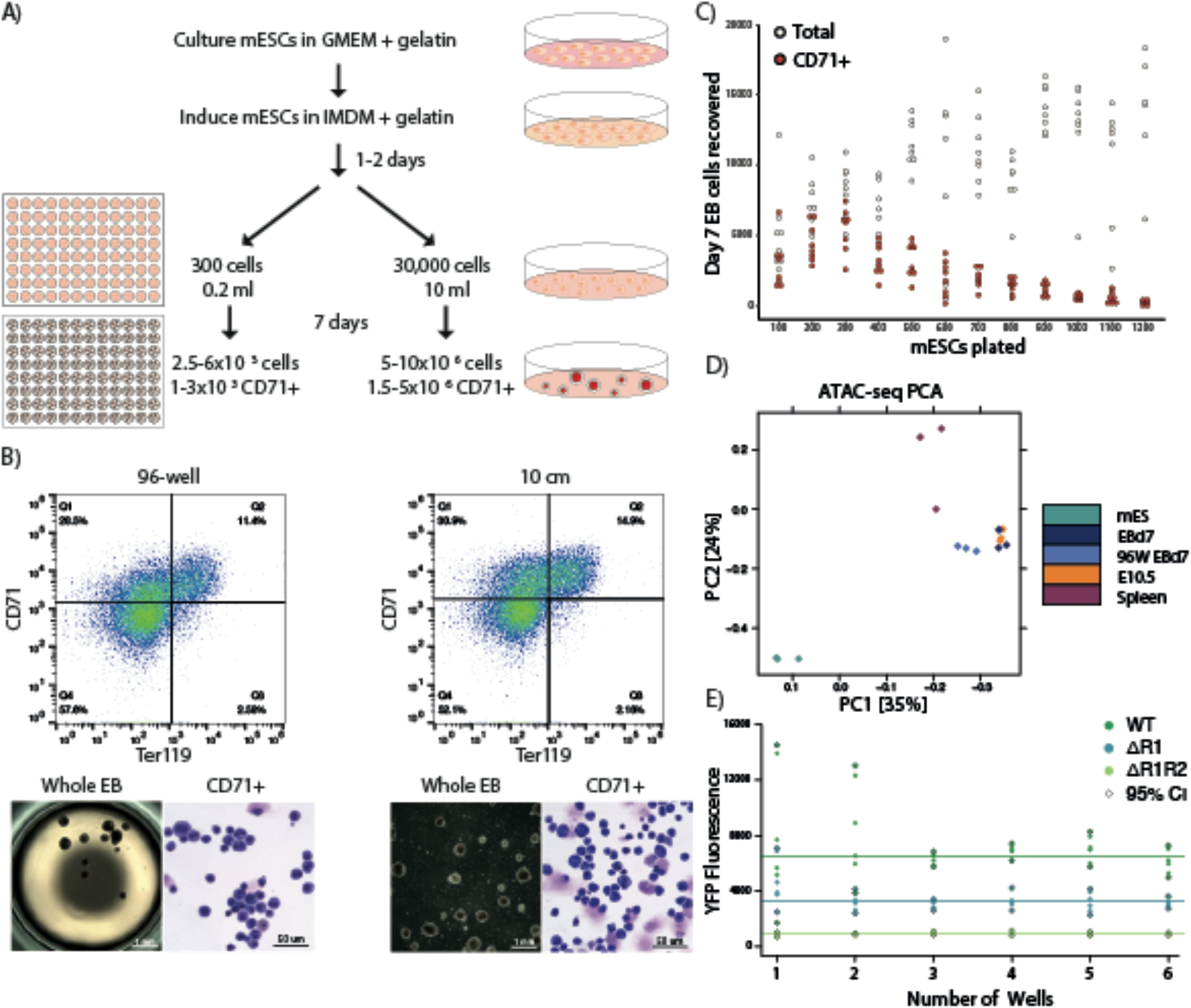
EB differentiation scaled down to a 96-well format. **A**) A schematic comparison of protocols for 10 cm-dish format and miniaturized 96-well plate EB differentiation. **B**) Characterization of the erythroid output from miniaturized EB differentiation by flow cytometry for CD71 and Ter119 erythroid markers, gross-scale microscopy of whole EBs and cellular MGG staining. Whole EB images show a single 96-well and a sample of a 10 cm dish, respectively. **C**) Total and CD71+ cell counts from day 7 EBs for a range of mESCs plating densities in 200 μl differentiation media in 96-well format. Optimal CD71+ output is at around 300 cells/well. **D**) PCA plot of genome-wide ATAC-seq peaks, as in Figure 2B, with the addition of CD71 + cells from 96-well EBs (96W d7EB). 96-well miniaturized EB culture clusters with bulk day 7 EB-derived cells (EBd7) and E10.5 embryonic erythrocytes away from mESCs and spleen-derived erythrocytes. **E**) Median YFP fluorescence readings from erythroid cells from 1-6 combined wells of a 96-well plate containing EBs for three genetically engineered α-globin YFP-tagged mESC lines: wildtype (WT), enhancer R1 knockout (ΔR1) and enhancers R1, R2 knockout (ΔR1R2). Solid lines indicate the average median YFP fluorescence for each cell line. Points highlighted with a black outline represent the 95% confidence interval (CI) values for each dataset. Combining 3 wells is enough to reduce output variability/genotype.

To further validate the equivalence of 96-well EB-derived erythroid cells to those from the routine 10 cm dish culture, we applied the column CD71+ purification to a pool of 96-well EBs and performed ATAC-seq. As for conventionally cultured EB-derived cells, the 96-well EB-derived isolated erythroid cells showed specific chromatin accessibility over the embryonic globin genes and clustered together with primitive cells by genome-wide analysis of ATAC-seq data (Figure 5D; Supplemental Figure 3B). Together, these results indicate that it is possible to scale down the haematopoietic EB differentiation procedure to a miniaturized 96-well format, without compromising the efficiency and faithfulness of differentiation.

Variability in EB size and haemoglobinization is common in both large- and small-scale cultures.^35^ The resulting impact on globin gene expression from single cells (as seen in Figure 3A) is generally averaged out when plating and analysing populations at a large scale, minimising any potential plate-to-plate experimental variability. However, the potential impact of well-to-well experimental variability could be severe when harvesting just a few EBs at a time, due to poor sampling of the erythroid populations. Although we optimized plating density conditions to maximize the CD71+ cell output per well, it was unclear whether this number of erythrocytes (~1-3×10^3^ CD71+ cells from 1-10 EBs per well) would be sufficient to provide a reliable readout of gene expression.

To that end, we studied the variation in α-globin-YFP fluorescence levels from EBs from three genetically modified mESC lines, each of which produces a different level of fluorescence (Figure 5E). The parental WT α-globin-YFP line (as used in Figure 1C) sets the maximum possible signal output from the locus; the ΔR1 line produces a reduced signal (~40% reduction); the ΔR1R2 line sets the lowest signal in the panel, reflecting the dramatic reduction (~90%) of alpha-globin expression seen in the absence of these two major enhancers.^21^ Using this panel to calibrate the system, we found that combining EBs from three wells (representing a of total ~12,000 CD71+ cells) was sufficient to lower the variation between readings; combining further wells beyond this number did not improve the data quality.

The ease of sample processing in a 96-well format makes this miniaturized format the optimal system when managing large numbers of clones in parallel. For more modest experimental designs, it is also possible to differentiate cells in a 24-well plate to reduce the need to combine multiple wells (Supplemental Figure 3D).

## DISCUSSION

The use of reliable and scalable *in vitro* cellular systems in which the cells produced have been well-characterised will be of great utility in facilitating research whilst reducing animal use and associated time and costs. In this report, we have addressed and developed solutions to four important issues when using mESCs to study haematopoiesis and in particular erythropoiesis. First, we have established a scalable, erythroid-cell specific purification protocol applicable directly to disaggregated day 7 EBs without the need for laborious replating steps. Second, we have defined the developmental origin of these spontaneously differentiated EB-derived erythroid cells; they uniformly represent primitive erythropoiesis. Third, we have shown that the purified erythroid cells faithfully recapitulate chromatin accessibility, three-dimensional folding and expression phenotypes of primary cells from both wildtype and manipulated genotypes. Finally, we miniaturized the process to a 24- and 96-well format, creating a protocol that is suitable for large scale screens. We, therefore, have harnessed the mESC-EB system as a simple and scalable source of well-characterized erythroid cells for genetic manipulation and molecular investigations.

The system we present here can be applied to generate mouse primitive erythroid cells for most existing experimental assays. By plating in the bulk 10 cm dish format, we provide a cost-effective, single-plating protocol to derive tens of millions of erythroid cells in a single experiment. The cell numbers obtained make samples amenable to a wide range of molecular biology techniques including assays for DNA accessibility (ATAC-seq; ~10^5^ cells), DNA binding events (ChIP-seq; 10^6^-10^7^ cells) and chromatin conformation assays (Capture-C; 10^4^ to 10^7^ cells). At the other end of the spectrum, the system can be used to multiplex the differentiation of hundreds of genetically modified clones for mid-throughput parallel analysis. These experiments would be unachievable using mouse models. High-throughput readouts such as flow cytometry for tracking an immunophenotype or a fluorescent tag are ideally suited to this system, and make the system an attractive option for genetic or chemical screens.

We have achieved such scalability in the system by optimizing culture conditions and validating the output. One of the challenges is the inherent heterogeneity of EB growth in bulk culture.^36,37^ This increases differentiation variability in the erythroid population and renders the system unreliably noisy. Our data are in line with previous findings about the nature of erythropoiesis within EBs and the observation that particularly dense cultures of EBs favour cardiac rather than haematopoietic differentiation.^38^ Successful differentiation of other lineages of all germ layers within EBs has also been found to be directly influenced by EB size.^39,41^ Alternative methods such as hanging drop differentiation^36,42^ promote more uniform EB growth but are technically more challenging and not amenable to high throughput assays. Alternatively, the culture of uniform single-sized EB in a single 96-well has also been developed.^43^ However, this is laborious and does not guarantee a uniform differentiation output. From our experience, the effect of EB variability is reduced in bulk cultures (10cm dishes). When we optimized for miniaturization, we maintained the same approach followed for bulk culture; plate an optimal cell number in culture dishes that allow EB aggregation, with minimal monitoring. We used the bulk culture and various reporter mESC lines to calibrate the output. Thus, the 96-well plating method we developed accounted for optimal cell numbers that reliably reported the phenotype whilst preserving the simple, cost-effective nature of the bulk culture.

Previous studies of erythropoiesis in the mESC-EB system focussed mostly on progenitor readouts in colony assays such as simple morphological assessments (colony shapes and cellular staining), limited immunophenotypic and RT-PCR evaluation of transcriptional programs. Previously, the EB differentiation system was mostly exploited to access and dissect rare populations that are otherwise inaccessible from *in vivo* models or to examine phenotypes precluded in a living organism.^44–47^ The ability to harvest scalable numbers of defined cells expands the range and depth of questions that can now be addressed using the mESC-EB system. Previous attempts to produce bulk erythroid cultures from EBs have used a laborious multi-step protocol of EB growth, disaggregation and plating of progenitors, requiring the use of expensive cytokine supplements and resulting in enucleated definitive red cell output of unclear origin.^48^ The spontaneously differentiated EB-derived erythroid cells described here provide an alternative simple and cost-effective approach. Although well-reported in the literature, these cells have not been well characterized. As EBs produce both primitive and definitive erythroid progenitors at various stages of their *in vitro* development, the spontaneously differentiated erythroid cells present at day 7 could be of both primitive and definitive origins. By analysing the chromatin environment and single-cell transcriptional output from the purified erythroid population we have shown that these cells uniformly resemble primitive erythroid cells.

The new protocol described here provides easy access to large numbers of well-defined mouse primitive erythroid cells. Together with the ever-expanding suite of genetic engineering and genomics tools, this work provides a robust and powerful system to address key, unanswered questions in molecular and cell biology using erythropoiesis as a model.

## ACKNOWLEGEMENT

This work was supported by the Medical Research Council (MRC Core Funding number, MR/T014067/1), H.S.F. Welcome Trust Studentship (109097/Z/15/Z), and R.A.B. Sir Henry Wellcome Fellowship (209181/Z/17/Z).

We would also like to thank Prof Merav Socolovsky and Samuel Wolock for their advice on the erythroid markers included in the Biomark experiment design.

## AUTHORSHIP CONTRIBUTIONS

M.T.K. conceived the project. H.S.F and M.T.K. designed experiments. H.S.F, C.L.H., R.A.B, A.J.K, M.E.G., D.M.J., C.B. and M.T.K. performed experiments. H.S.F, C.L.H., R.A.B, A.J.K, M.E.G., J.W.B., C.B. and M.T.K. analyzed data. H.S.F., R.A.B., A.J.K., and M.T.K. generated essential reagents. M.T.K. supervised the work. H.S.F and M.T.K. wrote the manuscript and made the figures with assistance from R.A.B and D.R.H and feedback from all the authors. Funding was acquired by D.R.H.

